# CODMA-A3: A Competitive Oligo-based DNA Modification Assay Applied to APOBEC Enzymes

**DOI:** 10.64898/2026.10.27.760525

**Authors:** Colin Rorex, Rachael Springman-Rodriguez, Armando Mendez, Jake Lehle, Elizabeth Sommer, Diako Ebrahimi

## Abstract

Quantifying the functional effects of human variation in proteins is a challenge because bioinformatic pipelines allow variant detection to outpace experimental characterization. This results in a practical problem of resource allocation to identify the most scientifically significant variants to prioritize for mechanistic investigation. There are many approaches appropriate depending on class of enzymatic activity being investigated. However, fundamental to many of these approaches is a trade-off between mechanistic insight and throughput.

In seeking to understand how activity changes across human variants of the APOBEC3 family of proteins we sought to create an early screening assay to prioritize variants for mechanistic study. We wanted an assay that was not dependent on protein purification, allowed for multiple-substrate competition and could be executed in a microtiter plate allowing for high throughput and approximate normalized substrate turnover without requiring classic Michaelis–Menten approaches.

Here we describe CODMA (Competitive Oligo-based DNA Modification Assay) and applied it to A3 enzymology as a method to rapidly screen A3 constructs for changes in substrate utilization fingerprints. CODMA-A3 detected effects of substrate competition, distinguished fluorophore identity from substrate sequence effects, resolved activity changes caused by a single nucleotide substitution within a competing substrate, and identified shifts in substrate utilization following modification of lysate composition, oligonucleotide architecture, or antibody-mediated protein perturbation.

CODMA-A3 occupies a methodological niche between low-throughput mechanistic assays and sequencing-intensive approaches by providing a rapid, inexpensive screening platform for comparative characterization of 20-40 enzyme variants prior to detailed mechanistic investigation.

**Graphical Abstract:** 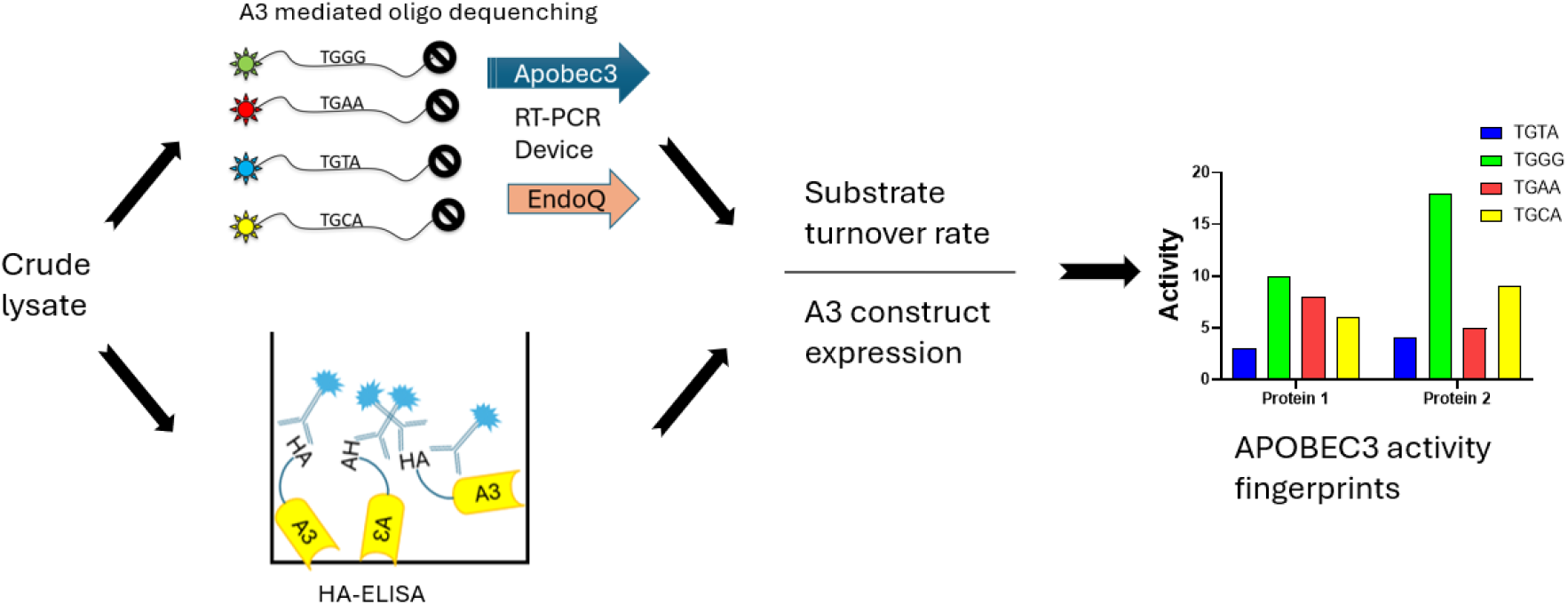

## Introduction

DNA-modifying enzymes generate diverse genomic outcomes that influence cellular development, transcriptional regulation, immune responses and oncogenesis^1, 2, 3^. When polymorphisms in DNA or RNA-modifying enzymes are identified or associated with a disease state, a fundamental question is how protein variations affect enzymatic function ^4^. An ideal method would allow quantification of enzyme kinetics in a competitive assay that better reflects the substrate competition encountered by DNA-modifying enzymes within cells while maintaining high throughput in a streamlined workflow^5,6^. However, existing approaches often involve protein purification, sequencing-intensive workflows, or interrogation of individual substrates in isolation. This means protocol selection often requires sacrificing throughput, multiplexing, or recovery of kinetic measurements in exchange for the others.

The APOBEC family comprises eleven cytidine deaminases that act on DNA and/or RNA substrates and participate in processes including mRNA editing, innate antiviral immunity, and antibody diversification^7^. Seven members of this protein family, known as APOBEC3 (A3) (A3A, A3B, A3C, A3D, A3F, A3G, and A3H) are cytosine deaminases that bind to single-stranded DNA and deaminate cytidine to uridine within distinct nucleotide motifs^8^. This catalytic activity can be detected by sequencing or strand breakage utilizing either uracil DNA glycosylase-NaOH dependent sequential reaction or direct endonuclease Q cleavage at the uridine base^9,10,11^. While these approaches generate high-quality measurements of deaminase activity, they generally rely on sequencing-based workflows or analysis of individual substrates in isolation, limiting their utility for rapid kinetic characterization of multiple enzyme variants across competing substrate contexts^9,10,12^.

Here we describe a Competitive Oligo-based DNA Modification Assay that detects A3 cytidine deamination by in-reaction endonuclease Q mediated strand breaks (CODMA-A3). Using quenched fluorescently labeled oligos allows for both multiplexing competitive substrates and detection of deaminase activity by increasing fluorescence. Samples can be run on a qRT-PCR platform such as the QuantStudio both to recover kinetic data and for on-machine deconvolution of oligo pools in a high throughput plate format. A paired anti-HA tag ELISA allows for quantification of A3 expression in 1xHA equivalents, enabling calculation of single-substrate turnover from a competitive pool. The resulting workflow can be executed in a single day with the ELISA and deaminase measurements performed in parallel by a single operator. This enables multiplex analysis of A3 substrate preferences and conversion of fluorescence signal to substrate turnover, allowing for calculation of apparent k_cat_ values directly from crude lysates. This workflow facilitates simultaneous characterization of multiple A3 proteins prior to more targeted mechanistic studies.

## Materials and Methods

### Expression of A3 constructs

A3 expression vectors in pcDNA3.1 HA vectors were obtained from Genscript using NCBI reference A3A/C/D/F/G/H CDS. The reference A3B CDS was modified with a chimeric intron (gtaagtatcaaggttacaagacaggtttaaggagaccaatagaaactgggcttgtcgagacagagaagactcttg-cgtttctgataggcacctattggtcttactgacatccactttgcctttctctccacag) at intron 5 to overcome toxicity in bacterial cultures. HEK293T cells were transiently transfected using PolyJet (SignaGen Laboratories, Frederick, MD USA, Cat# Sl100688) as per the manufacturer’s protocol for 6-well plate format transfections in duplicate. Culture media was replaced 24 hours post-transfection and harvested 48 hours post-transfection by mechanical disruption. Prior to harvest cells were washed in PBS, then overlaid with 500 μL of lysis buffer (25mM HEPES pH 7.4, 15mM EDTA, 10% glycerol and protease inhibitor, cOmplete Mini, Roche, St. Louis, MO USA Cat# 04693159001), harvested and pooled. An equal number of untransfected HEK293T cells were harvested to serve as producer controls for background subtraction of endogenous A3. Cells were lysed by a freeze-thaw cycle at - 80°C, then sonicated in a water bath sonicator (4°C, 40KHz, 100% power, sweeping) for 20 minutes in two 10-minute intervals. Lysed samples were clarified at 16,000g for 10 minutes and supernatants were aliquoted at 250 μL in fresh tubes and stored at -80°C. Purified human A3A protein was a generous gift from the laboratory of Dr. Reuben S. Harris (University of Texas Health San Antonio). The protein was supplied at a reported concentration of 63.7 μM. One microliter of stock protein was added to a 60 μL reaction master mix, corresponding to 21.2 pmol APOBEC3A per 20 μL reaction.

### HA-tagged protein quantification ELISA

A3 constructs were quantified by indirect ELISA run at room temperature with plate sealers to minimize evaporation until TMB substrate was added. The Pierce HA-tagged Positive Control (Thermo Fisher Scientific, Waltham, MA USA, Cat# 26180X) was diluted from the 1 μg/μL stock to a working 1 ng/μL stock, which was then used to generate a six point two-fold serial dilution series. Loading 10 μL of each standard per well results in a standard range of 10 ng-0.3125 ng. Samples here were run at 1 μL/well. In brief, standards and samples were immobilized by pipetting into medium binding polystyrene ELISA plates with 100 μL binding buffer (44 mM sodium bicarbonate, 11 mM sodium chloride, pH 9.6) in duplicate for 1 hour. Following binding, wells were washed three times with 300 μL of wash buffer (TBS + 0.05% Tween 20 v/v). Reagent buffer used for both blocking and antibody dilution was made by adding 1% BSA w/v to an aliquot of wash buffer. 100 μL of reagent buffer was added for 60 minutes to block non-specific reactions then washed. 100 μL of a 1:1000 dilution anti-HA detection antibody (Biolegend, San Diego, CA USA, Cat# 901513, RRID: AB_2565335) was added and incubated for 1 hour followed another wash. 100 μL of a 1:1000 goat anti-mouse HRP antibody was added and incubated (Invitrogen, Carlsbad, CA, USA, Cat# 31430, RRID: AB_228307) for 1 hour and washed. 50 μL of TMB substrate (Invitrogen, 00-4201-56) was then added and allowed to develop for 30 minutes before adding 50 μL 1M H_2_SO_4_. Plates were read at 450nm.

Quantification of samples was determined by interpolation from the six-point standard curve after background subtraction (Figure 1). 1x HA nmol equivalents were estimated by dividing the resulting mass (in ng) by 1083 ng/nmol (estimated mass of 1x HA peptide sequence).

**Figure 1:**
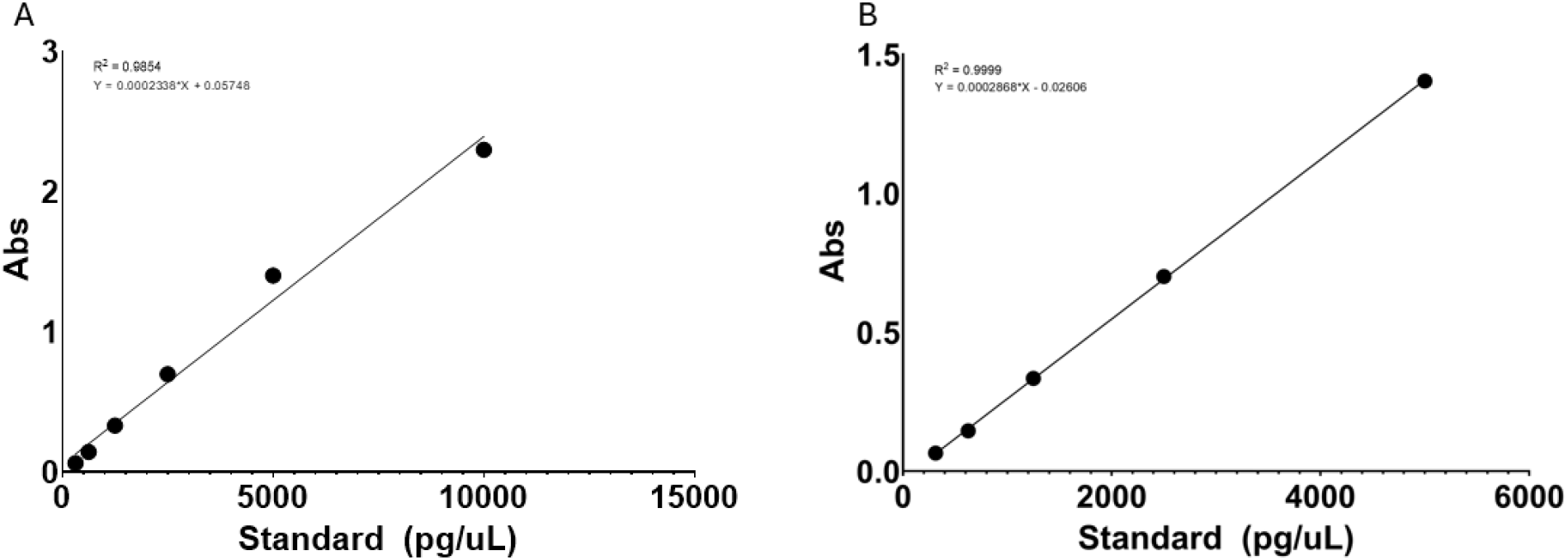
Standard curve generated using 10 ng to 312.5 pg 1×HA peptide to quantify A3 expression. Linear regression including all standards (A, R^2^ = 0.9864). Linear regression of the 5 ng to 312.5 pg working range used for sample interpolation (B, R^2^ = 0.9999).

### Substrate oligos

Oligos were purchased from Integrated DNA Technologies (IDT, Coralville IA USA) (Table 1) as lab ready 100 μM, HPLC purified stocks. Several working pooled stock master mixes (mm) were made by diluting individual oligos to a 10 μM concentration in the same solution. Standards to interpolate RFU to pmol of substrate turnover were sample oligos lacking 3’ quenchers and function per channel independent of oligo sequence basis: TGGG-**FAM**, TGAA-**SUN**, TGCA-**TAMRA**, TGTA-**ROX**. The 5-oligo mm used for plate controls always contained TGGG-FAM, TGAA-SUN, TGCA-TAMRA, TGTA-ROX, EndoQ control-Cy5. Sample 4-oligo mm varied in composition from the following pool: TGGG-FAM, TGGT-FAM, TGAA-SUN, TGAA-ROX, TGCA-TAMRA, TGTA-ROX, TGTA-SUN and TGAA-44-SUN. Single oligo or restricted oligo pools were run at the same 10 μM per oligo concentrations.

**Table 1.**
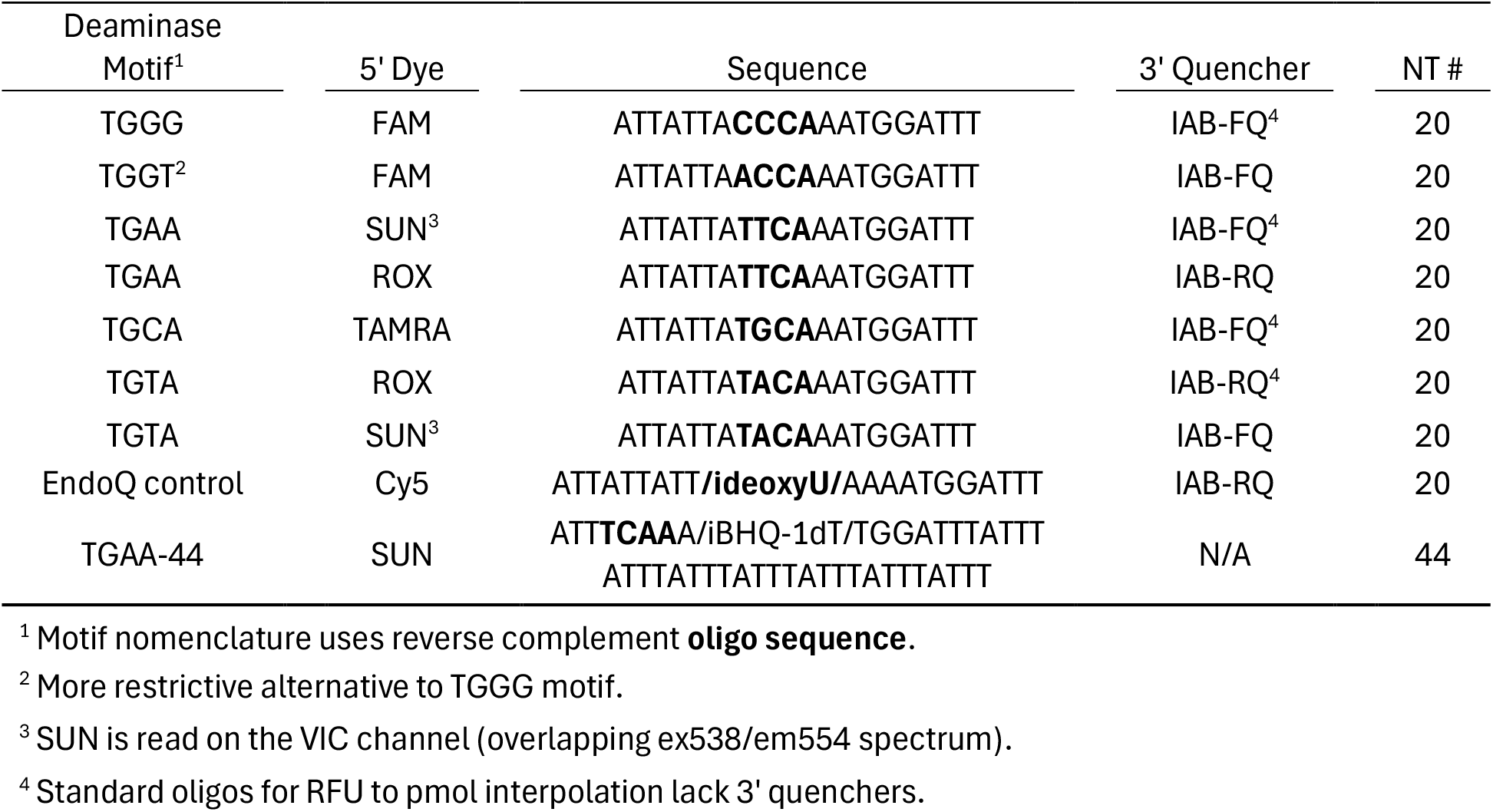
Apobec3 substrate oligos.

### CODMA-A3 protocol

Controls, samples and standards were prepared as described in Tables 2 and 3 and held on ice before being transferred to an optical 96 well plate (Bio-Rad Laboratories, Hercules, CA USA, Cat# HSS9641) and sealed immediately before running on a QuantStudio 6 Real-Time PCR System (Applied Biosystems, Thermo Fisher Scientific, Waltham, MA, USA).

**Table 2.** Standard for RFU to pmol interpolation^1^ (TGGG-FAM, TGAA-SUN, TGCA-TAMRA, TGTA-ROX)

| Standard | Source | Source vol | DEPC H<sub>2</sub>O | Final vol | per well |
| --- | --- | --- | --- | --- | --- |
| 10pmol | 10μM mm | 6.6 | 125.4 | 66 | 20 |
| 5pmol | 10pmol | 66 | 66 | 66 | 20 |
| 2.5pmol | 5pmol | 66 | 66 | 66 | 20 |
| 1.25pmol | 2.5pmol | 66 | 66 | 66 | 20 |
| 0.625pmol | 1.25pmol | 66 | 66 | 66 | 20 |
| 0.3125pmol | 0.625pmol | 66 | 66 | 132 | 20 |

<sup>1</sup>Volumes are expressed in μL.

**Table 3.** Control and sample preparation per well in μL.

| Sample | A3 vol<sup>3</sup> | EndoQ<sup>4</sup> | r2.1 buffer<sup>2</sup> | 4oligo mm<sup>5</sup> | 5 oligo mm<sup>5</sup> | H<sub>2</sub>O Bal<sup>1</sup> |
| --- | --- | --- | --- | --- | --- | --- |
| Oligo only control | NA | NA | 2 | NA | 1 | 17 |
| EndoQ control | NA | 2 | 2 | NA | 1 | 15 |
| A3/293T samples | 7 | 2 | 2 | 1 | NA | 8 |

<sup>1-5</sup>Superscript numbers indicate pipetting order.

Thermostable Endonuclease Q (EndoQ) and r2.1 buffer was obtained from New England Biolabs (NEB, Ipswich MA, USA Cat #M0701S). Plate controls (oligo only and EndoQ control) were run as single well reactions while all other samples and standards were run in technical triplicate. Data acquisition was set for FAM/ROX/TAMRA/VIC/Cy5 channels for every loaded well then reactions were incubated at 37°C for 15 s, followed by fluorescence acquisition every 3 min at 37°C for 40 cycles. Data export at the end of the run included multicomponent data.

### Modification of sample lysates

RNA depletion was performed directly on crude lysate samples before addition of oligo by replacing 1 μL H_2_0 volume with 1 μL of RNase solution (Qiagen, Germantown, MD, USA, Cat #19101) as prepared per the manufacturer’s instructions for body fluid purification from the PureGene protocol (Cat #158023). Both RNase A-treated and corresponding untreated samples were incubated at RT for 30 minutes prior to addition of 4-oligo substrate mix.

Anti-HA mediated antibody aggregation was done by replacing 6 μL of the H_2_0 volume prior to addition of 4-oligo mm with an antibody cocktail. First 3μL (1 μL per reaction) of a 1:1000 dilution in 1x TBS of primary ELISA antibody was added and incubated for 10 minutes at RT.

Next 3 μL (1 μL per reaction) of a 1:1000 dilution in 1x TBS of secondary ELISA antibody was added and incubated on ice for 15 minutes.

Purified protein samples replaced the entire volume of crude lysate with a corresponding increase in H_2_0 to maintain a 20 μL reaction volume per sample.

### Data analysis

Multicomponent data was segmented by fluorophore and treatment conditions utilizing a Python script (GitHub - github.com/ColinRorex/CODMA), which generated an Excel workbook for downstream analysis. Subsequent calculations including RFU to pmol interpolations, slope analysis and k_cat_ were made in Excel (Version 2607, Microsoft, Redmond, WA USA). EndonucleaseQ activity can be calculated by fitting Cy5 signal to a one phase association nonlinear curve using nonlinear least-squares regression. Oligo stability was validated by plotting fluorescence over time from oligo-only controls. To interpolate substrate turnover from RFU measurements, fluorophore standards were generated by averaging stable fluorescence (typically timepoints 15-20 to 40). Linear regression, log-transformed linear regression, and second-order polynomial regression models were evaluated. The model yielding the lowest residual sum of squares across the working dynamic range was used for interpolation of RFU measurements to pmol substrate concentrations. Substrate conversion rate was calculated either from primary RFU data or from RFU to pmol interpolation over a rolling 5 time point (15 minute) window using built in (Slope) and (RSQ) functions Excel. Background subtraction for endogenous A3 expression in HEK293T, reaction solution, and plate variability were done by subtracting the untransfected HEK293T values from the A3 construct samples on a fluorophore-by-fluorophore basis. Normalized activity was calculated by dividing background-subtracted reaction rates by the calculated 1×HA molar equivalents present in each reaction. All curve fitting, regression analyses, and interpolation procedures were performed in GraphPad Prism version 10 (GraphPad Software, Boston, MA, USA), while substrate kinetics were calculated in Excel. Statistical analysis was performed using GraphPad Prism as indicated in text based on number of comparisons with pairwise comparisons done with Tukey’s post hoc pair-wise comparison.

Because enzyme abundance is estimated from ELISA-derived HA-tag equivalents and substrate turnover is calculated from interpolated fluorescence measurements at fixed concentrations and not by Michaelis-Menten kinetic analyses, all reported k_cat_ values should be interpreted as apparent kinetic parameters suitable for comparative analyses across A3 proteins, substrates, and experimental conditions rather than as absolute catalytic constants.

## Results

### Quantification of A3 expression by HA-tag ELISA

Calculation of substrate turnover (k_cat_) requires quantification of enzyme concentration within the reaction. As crude cellular lysates are employed in this method, we utilized an indirect ELISA technique rather than a tag-based purification approach to facilitate high throughput processing and to maintain sample composition identical to that used in the CODMA-A3. A commercial kit can be utilized but we used a custom direct ELISA with a previously validated anti-HA antibody and HRP-conjugated secondary. A standard curve spanning 10 ng to 312.5 pg HA peptide standards provided a suitable dynamic range for interpolation of A3 expression as 1× HA molar equivalents from 1 μL of lysate (Figure 1, Table 4).

**Table 4.** Estimated A3 protein expression.

| A3 | Accession | pmol/uL |
| --- | --- | --- |
| A3A | NM_145699.4 | 6.6142 |
| A3B | NM_004900.5<sup>1</sup> | 1.9772 |
| A3C | NM_014508.3 | 3.7796 |
| A3D | NM_152426.4 | 1.0829 |
| A3F | NM_145298.6 | 0.5377 |
| A3G | NM_021822.4 | 1.8071 |
| A3H | 183 - NM_181773.5 | 3.9191 |
|  | 200 - NM_001166003.3 | 1.0905 |

<sup>1</sup> Contains a modified intron 5 sequence.

### In-plate controls to recover assay-specific signal

To verify the CODMA-A3 performance and ensure the results are not a product of non-specific reactions, each experimental run must include an oligo only control (Figure 2a, Table 3) to test for oligo dequenching, an Endonuclease Q control (Figure 2b, Table 3) to verify cleavage specificity for deaminated cytosines and adequate Endonuclease Q enzymatic activity. The HEK293T producer control (Figure 2c, Table 3) serves as background correction for plate optics, sample composition, and endogenous expression of A3 proteins in HEK293T cells. The producer control must be conducted in technical triplicate then averaged on a per-substrate basis before being subtracted from sample wells for each corresponding fluorescent channel.

**Figure 2.**
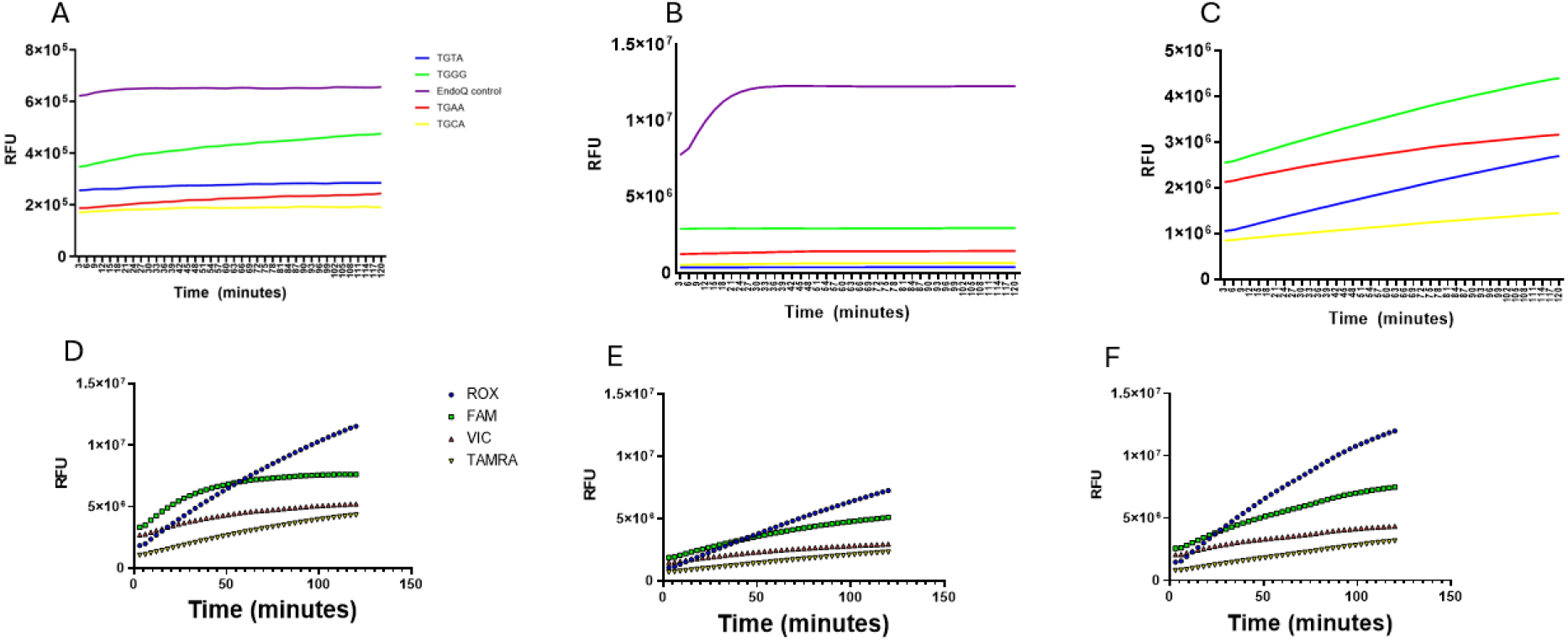
In plate standards and controls for oligo stability, Endonucease Q-specific activity and producer HEK293T background deaminase activity. Oligo-only control reactions demonstrate fluorophore and quencher stability throughout the assay (A). Increased Cy5 fluorescence in the Endonuclease Q control reaction demonstrates substrate-specific cleavage activity and rapid substrate exhaustion, while non-target fluorophore channels remain unchanged (B). Deaminase activity of endogenously produced A3 proteins in HEK293T producers can be detected and quantified (C). Averaged RFU values from lysates transfected with plasmid vectors expressing A3A (D), A3F (E), and A3H (F) demonstrate unprocessed substrate turnover.

### Interpolation preserves substrate-specific kinetic relationships observed by fluorescent readouts

The primary readout of CODMA-A3 is fluorescent evolution, a direct readout of quencher cleavage without additional calculation (Figure 2d-f). A secondary derived readout of estimated per-substrate turnover can be calculated by using a 6-point standard curve (Table 3). Here in both RFU and pmol space several time intervals were considered for rate calculations, and we found high degrees of linearity out to a 5-point (15 minute) rolling window across multiple operators and independent experiments before background subtraction (Figure 3a&b, representative data for A3F). Standard ranges used for interpolation of RFU measurements to substrate turnover (pmol) are anchored in substrate pool design, with the upper limit corresponding to 100% cleavage of 10 pmol of quenched oligos. This precludes utilizing 4PL fit as top-end saturation fitting would both require the expansion of the standard range and necessitate running both single channel and multiplex standards followed by manual deconvolution methods. Expansion of the standards will also shift the assay to a 384 well format. Comparison of raw fluorescence and substrate turnover rates demonstrates that substrate-specific enzymatic fingerprints are preserved following standard interpolation (Figure 3c).

**Figure 3.**
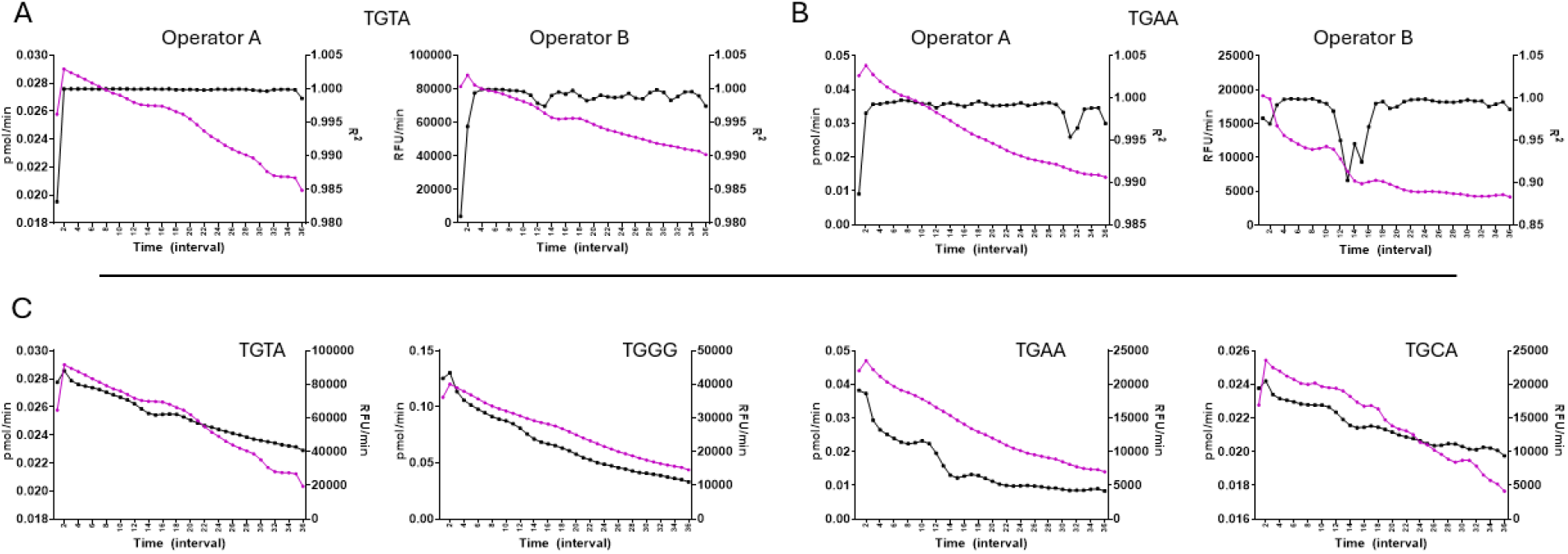
Rolling-window analysis demonstrates robust kinetic estimation across operators and processing workflows, while interpolation of substrate turnover preserves assay kinetics. A3F assay data from two independently executed experiments showing interpolated pmol/min or RFU/min (black, square, primary axis) together with linearity of 15-minute rolling (purple, round secondary axis) for the single substrates TGTA (A) and TGAA (B). Comparison of decay rates derived from pmol interpolation (purple, round, primary axis) and primary RFU measurements (black, square, secondary axis) demonstrates similar kinetic trajectories across all substrate motifs (C). Data represent averaged triplicates and interval denotes for 15-minute rolling window where interval 1 corresponds to 3-15 minutes and 2 corresponds to 5-18 minutes.

### Interpolation of substrate turnover allows quantitative comparison of A3 activity across proteins and substrates

Replicate experiments of reference A3A-H demonstrate that raw fluorescence data can be used to recover substrate-specific activity profiles across multiple A3 proteins and compare rates on a per-oligo basis (Figure 4a). However, interpolation of RFU measurements to pmol substrate turnover followed by normalization to A3 abundance from ELISA-derived quantifications enables calculation of apparent k_cat_ values, facilitating direct comparison of enzymatic activity across both substrates and A3 family members (Figure 4b).

**Figure 4.**
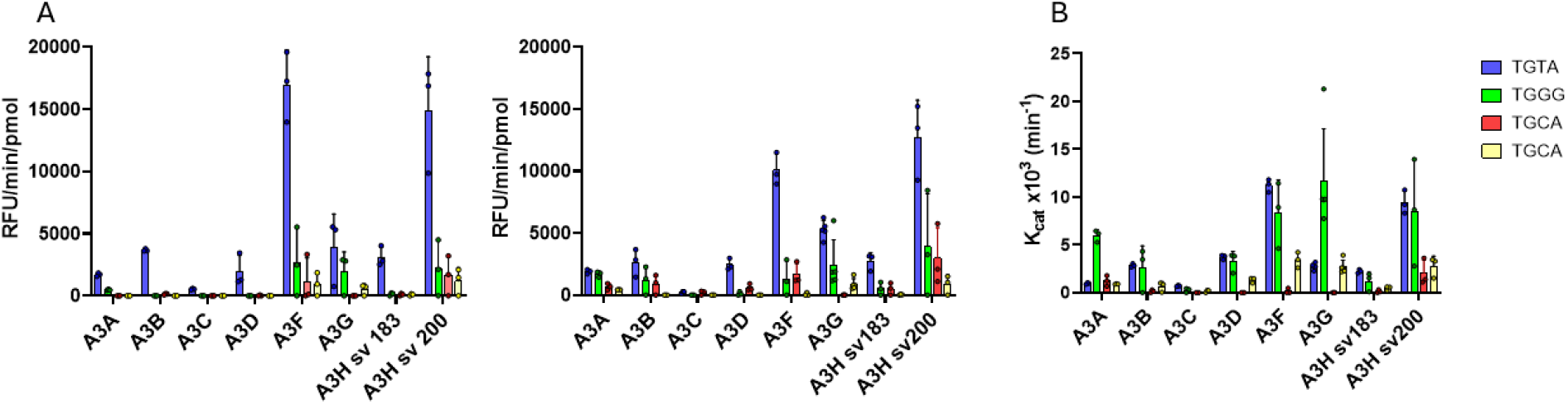
Interpolation of fluorophore-specific standard curves enables quantitative comparison of substrate turnover across independent targets. Replicate experiments of reference A3A-H with competitive substrate pools demonstrate similar oligo preferences (A). Interpolation of substrate turnover into pmol enables quantification of early reaction kinetics and correction for fluorophore-dependent signal effects, allowing calculation of apparent k_cat_ values (B). All data are background-subtracted and normalized to A3 expression.

### CODMA-A3 resolves substrate-dependent differences in APOBEC activity

We next asked whether CODMA-A3 could resolve biologically meaningful aspects of substrate context. Specifically, we examined the effects of substrate competition, fluorophore selection, and subtle sequence variation within competitive substrate pools. First, we tested whether pooled oligo substrates exhibited substrate competition or simply reflected the summed activity observed in individual substrate reactions. To address this, 4-oligo pools were run alongside equal-molar single-substrate reactions for A3G and A3H within the same experimental plate (Figure 5A). Next, we tested whether the fluorophore attached to a specific oligo motif influenced substrate turnover rates. The standard substrate pool was modified by swapping the fluorophores assigned to the TGAA and TGTA oligos, generating TGAA-ROX and TGTA-SUN while maintaining identical oligo sequences. We did not find a significant change in substrate turnover for either A3A (TGTA-ROX 0.96 ± 0.08, TGTA-SUN 1.56 ± 0.836, p = 0.443; TGAA-SUN 1.23 ± 0.56, TGAA-ROX 1.09 ± 0.63, p = 0.951) or A3H sv200 (TGTA-ROX 9.42 ± 1.22, TGTA-SUN 10.97 ± 3.42, p = 0.592; TGAA-SUN 2.04 ± 1.37, TGAA-ROX 5.89 ± 0.5, p = 0.083), and the substrate rank-order preferences remained unchanged (Figure 5b), indicating that substrate preference measurements are driven by oligo sequence rather than fluorophore identity. Finally, we tested the sensitivity of CODMA-A3 to subtle changes in substrate motif context. The TGGG substrate was replaced with a TGGT substrate in an otherwise identical competitive substrate pool and assayed in A3G and A3H SV200 lysates to quantify substrate turnover (Table 1). TGGT oligo pool data was combined with previous experimental data of TGGG oligo pool A3G and A3H SV200 allowing analysis by two-way ANOVA using protein and substrate pool comparisons as factors (Figure 5c). Despite differing by a single nucleotide, replacement of TGGG with TGGT altered the recovered substrate preference profiles (A3G: TGGG 33.9 ±16.99, TGGT 15.33 ± 4.73, *p* = 0.0169; A3H SV200: TGGG 25.36 ± 16.76, TGGT 50.55 ± 19.24, *p* = 0.0023). In contrast, activity measurements for the remaining shared substrates were largely preserved between experiments, indicating that CODMA-A3 is sensitive to subtle sequence-dependent effects while maintaining consistent substrate-specific activity measurements across independent competitive pools. These results demonstrate that CODMA-A3 can resolve subtle sequence-dependent differences in substrate selection within a competitive assay environment.

**Figure 5.**
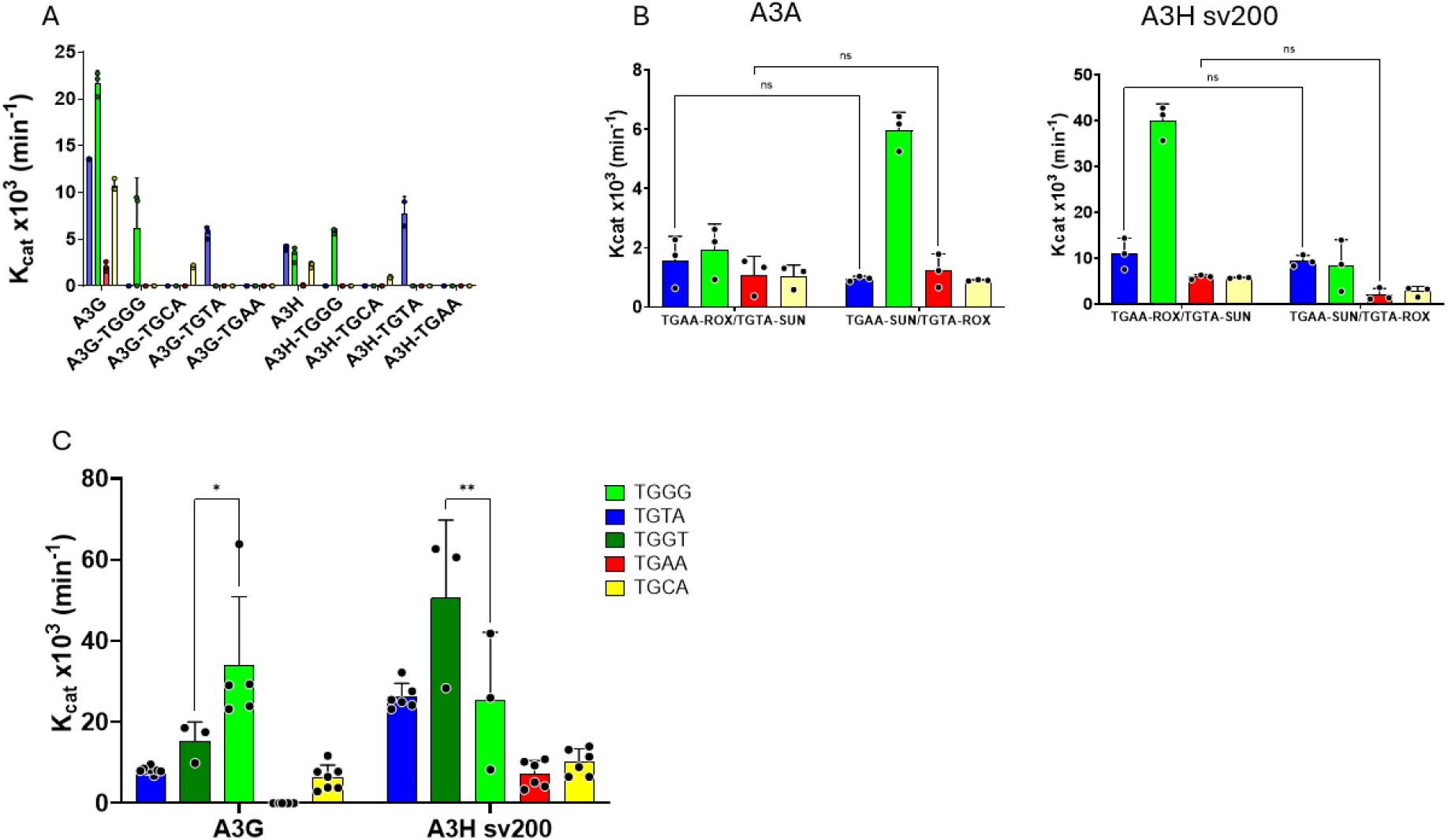
CODMA-A3 resolves biologically meaningful aspects of substrate context. Multiplexed and decomposed single-oligo reactions for A3G and A3H sv183 show competition is not predicted by the sum of single substrate activity (A). Changing to alternative fluorophores for TGAA (SUN to ROX) and TGTA (ROX to SUN) does not substantially alter oligo-specific substrate turnover rates or substrate preference hierarchies (B). Replacement of the TGGG motif oligo with a more restrictive TGGT alters A3-specific preferences. Data shown are from two independent experiments plotted together (C). ns *p* > 0.05, * *p* < 0.05, ** *p* <0.01 using two-way ANOVA with Tukey’s correction.

### Manipulation of lysate composition is detectable by CODMA-A3

We next sought to determine whether CODMA-A3 could tolerate experimental manipulations while maintaining protein- and substrate-specific A3 activity. To test this, we evaluated three distinct modifications to the assay framework: perturbation of A3 lysate composition, alteration of substrate architecture, and antibody-mediated perturbation of the target enzyme.

Perturbation of A3 lysate composition - We incorporated both RNase A pretreatment of lysate and purified A3A to test CODMA-A3 chemistry tolerance to sample modification and whether activity profiles of crude lysate reflect purified A3 activity. Here we found that substrate rank order hierarchies were preserved with only TGGG substrate turnover showing a difference in turnover under conditions of RNase A treatment (RNase A 3.04 ± 3.4, crude lysate 5.96 ± 0.62, purified 3.80 ± 2.24, RNase A vs crude lysate *p* = 0.020, purified vs crude lysate *p* = 0.43) (Figure 6a).

**Figure 6.**
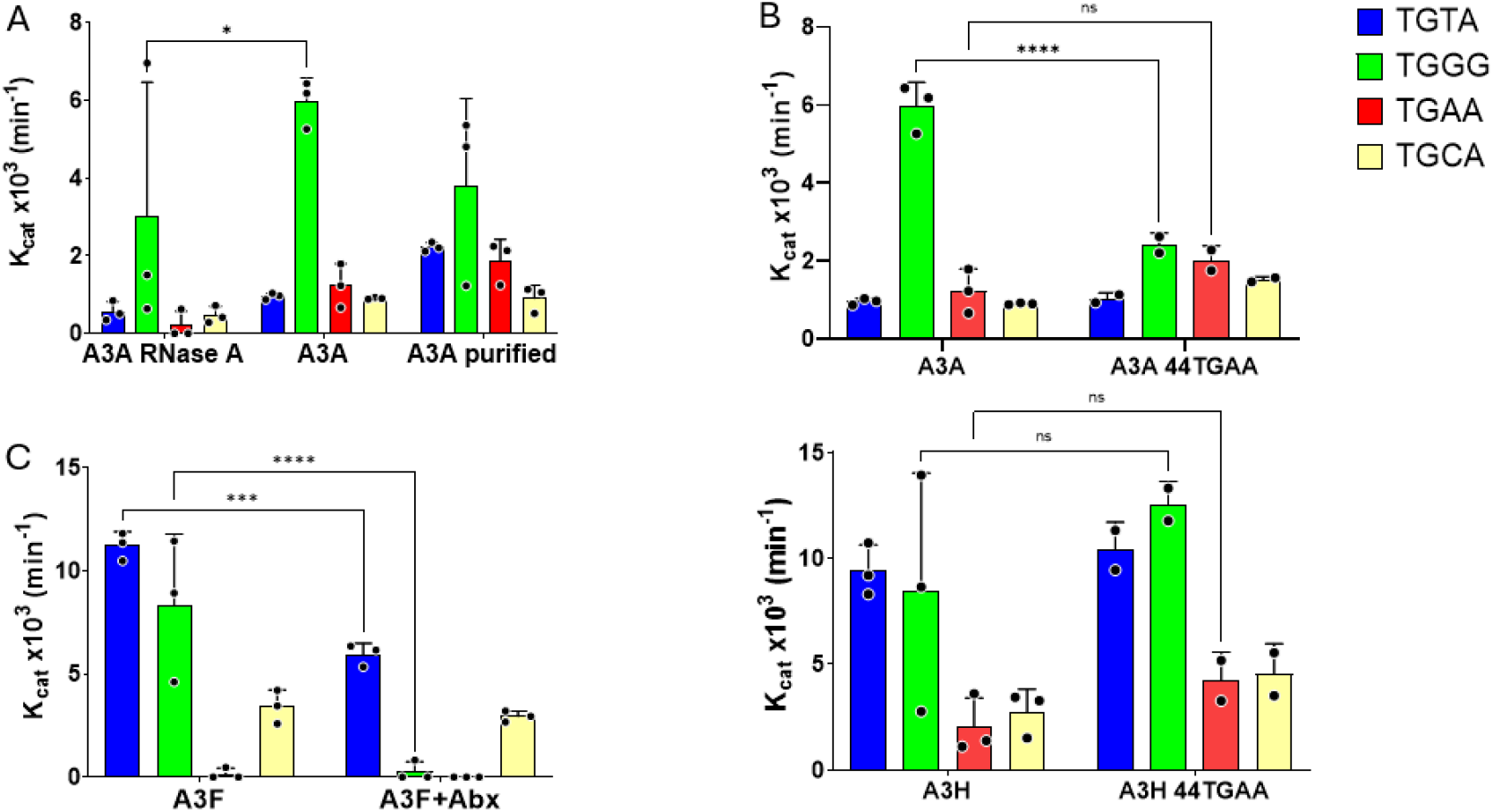
CODMA-A3 assay can detect shifts in substrate turnover under conditions of lysate modification. Normalized substrate turnover data from A3A crude lysates pretreated with RNAse A (left) untreated (middle) or purified A3A (right) show preserved substrate preference hierarchies (A). Extension of the TGAA oligo from a 20 to 44 nucleotides does not affect deamination rate (B). Pre-treatment with primary and secondary antibodies utilized in quantification ELISA show a substrate-specific reduction in deaminase activity (C). ns *p* > 0.05, *** *p* < 0.001, **** *p* <0.0001 using two-way ANOVA with Tukey’s correction.

Alteration of substrate (oligo) architecture - Next, we tested whether CODMA-A3 could detect reported effects of oligo length on A3 activity by designing a 44nt TGAA-SUN oligo (Table 2) and replacing it in the substrate mix while holding the other oligos constant as a minimal modification to the oligo pool. While we found no change in substrate turnover for the TGAA oligo in A3A (20nt TGAA 1.23 ± 0.563 vs 44nt TGAA 2.02 ± 0.37, *p* = 0.15) or A3H (20nt TGAA 2.04 ± 1.38 vs 44nt TGAA 4.23 ± 1.35, *p* = 0.84), we did find that TGGG substrate turnover decreased in the A3A sample (TGGG – 20nt TGAA 5.96 ± 0.62 vs TGGG – 44nt TGAA 2.42 ± 0.30, *p* < 0.0001) but not the A3H sample (TGGG – 20nt TGAA 8.45 ± 5.59, TGGG – 44nt TGAA 12.56 ± 1.09, *p* = 0.36) (Figure 6b). This A3A-specific decrease in TGGG turnover may suggest a shift in oligo preference.

Antibody-mediated perturbation of A3 - Finally, we tested whether antibody-mediated protein aggregations could perturb A3 activity measured by shifts in substrate fingerprint. We found that when adding the anti-HA primary and secondary antibodies used in the ELISA quantification, A3F showed a near complete ablation of TGGG turnover (untreated 8.33 ± 3.46 vs Abx treatment 0.28 ± 0.48, *p* < 0.0001) but only around two-fold reduction in TGTA turnover (untreated 11.2 ± 0.68 vs Abx treatment 5.96 ± 0.54, *p* = 0.0006) (Figure 6c). This result suggests that antibody aggregation is affecting a fundamental component of A3F mechanistic activity. This demonstrates that CODMA-A3 can detect the motif-specific consequences of antibody-mediated perturbation in crude lysates.

Collectively, these results highlight that CODMA-A3 as a screening assay that combines flexibility in assay design with high sensitivity for generating biologically meaningful data in a high throughput workflow.

## Discussion

Characterizing functional effects of protein variation is challenging because observed variant activity reflects the combination of substrate preference, retention, release and catalytic activity. The experiments to characterize individual contributions have practical tradeoffs where depth of biological insight comes at the expense of speed and resources. When bioinformatic analysis identifies multiple candidate variants this investment of time and resources required for comprehensive characterization accumulates rapidly. This necessitates either a large expenditure of resources to characterize them all or some form of priority triage.

Investigating A3 function benefits from extensive prior work elucidating both mechanism and substrate preference, resulting in a range of experimental approaches. Single-substrate gel-based endpoint cleavage assays allow for sequential interrogation of preferred substrate motifs^11^. Modifying substrates with quenched fluorophores allows for kinetic measurement but requires prior purification^9^. Sequencing intensive methods such as UDP-seq excel at mapping sequence preferences across complex libraries or genomic contexts, generating endpoint data that require substantial downstream processing^10^. Common to many of these workflows are protein purification or single substrate reactions, which decouple A3 activity from the competitive multi-substrate cellular environment.

CODMA-A3 was constructed to reduce these constraints simultaneously. The assay places multiple fluorophore-labeled, quenched oligonucleotides that carry distinct dinucleotide contexts into a single reaction volume, allowing substrate competition itself to become the central experimental variable. Continuous multi-channel fluorescence on a standard qPCR instrument reports the real-time fate of each competitor without sequential reactions or library preparation. Parallel ELISA quantification of HA-tagged enzyme abundance converts fluorescence rates into apparent catalytic constants, enabling direct numerical comparison of activity across family members, variants, and experimental conditions from crude lysates. The workflow can be completed in a single day by a single operator, recovering multi-substrate kinetic fingerprints under conditions that more closely approximate cellular substrate competition than pure single-substrate measurements.

In investigating the sensitivity of CODMA-A3, we found that multiplexed substrate profiles are not simple additive combinations of single-substrate rates, consistent with substrate competition. Substrate utilization was driven by enzyme-substrate interactions rather than fluorophore identity, as demonstrated by exchanging fluorophores between substrates. Single-nucleotide substitution within an otherwise shared sequence context revealed protein-specific shifts in utilization, demonstrating that the assay possesses sufficient resolution to detect biologically meaningful sequence differences. Experimental manipulations of lysate composition (RNase pretreatment and/or enzyme purification), oligonucleotide architecture (length variation), and antibody-mediated aggregation are readily detectable by multi-substrate fingerprint. These results indicate that CODMA-A3 can function as an upstream screening platform capable of identifying conditions or perturbations that merit more resource-intensive mechanistic follow-up.

During development of CODMA-A3, the sensitivity and modular design of the assay revealed three immediate extension opportunities. First, the ability to quantify endogenous HEK293T A3 activity enables CODMA to operate in any other cell line or primary cell type with equal or greater A3 activity. PBMCs have high A3 expression that is exacerbated by inflammation and should be highly compatible with CODMA^13,14^. Second, CODMA fundamentally reports DNA modification as fluorescence by cleavage of quenched oligos. There are many classes of DNA modifications that can be paired with restriction endonucleases. Here DNA methyltransferases can be coupled to MspJI and HpaII. This opens up CODMA as a platform for other DNA modifications not just A3 cytidine deaminases. Third, while CODMA as designed operates by detecting increased fluorescence, a qRT-PCR platform can also detect decreases in fluorescence. Here the assay would work in reverse utilizing two shorter oligos, one with a fluorophore, the other with a quencher in conjunction with a reverse complement scaffold for ligation. This would result in quenching of the reporter, allowing CODMA to interrogate DNA repair enzymes.

CODMA-A3 occupies a methodological position between high-resolution single-substrate kinetics and high-complexity sequencing surveys of DNA-modifying enzymes. It was developed to fill a very specific niche as a method for rapidly screening 20-40 protein constructs to identify the most scientifically exciting variants for deeper mechanistic investigation. Bioinformatics and deep-sequencing datasets allow for variant discovery faster than experimental characterization. By developing CODMA-A3 to operate in crude lysates, we avoid the purification bottleneck. In practice, we found that parallel transient transfections for variant expression were our largest bottleneck. CODMA-A3 has a very low infrastructure requirement, requiring access to a qRT-PCR platform enabling kinetic and competitive substrate interrogation. Importantly, this assay supports manipulation of both substrate and sample composition, becoming a low-cost framework for hypothesis testing.

## Contributions

**Colin Rorex:** Conceptualization, Methodology, Software, Validation, Formal Analysis, Investigation, Data Curation, Writing – Original Draft and Editing, and Visualization. **Rachael Springman-Rodriguez:** Conceptualization, and Validation. **Armando Mendez:** Investigation, and Writing – Review and Editing. **Jake Lehle:** Methodology and Writing – Review and Editing. **Elizabeth Sommer:** Validation. **Diako Ebrahimi:** Conceptualization, Supervision, Funding acquisition, Writing – Review and Editing.

## Declaration of Generative AI Use

Generative artificial intelligence (AI) tools based on large language models (LLMs) were used to assist in the development and debugging of Python scripts for raw data segmentation, and to support language editing of manuscript drafts. All code, analyses, results, interpretations, and manuscript content were reviewed and verified by the authors, who take full responsibility for the final work.

## Acknowledgements

That work was funded by the NIH/National Institute of Allergy and Infectious Diseases (NIAID) grant for HIV research (R01AI179465-01A1 D.E.). Purified A3A utilized in this project was the kind gift from the lab of Dr. Reubin S. Harris of UT Health San Antonio.

